# Rapid Fractionation and Characterisation of Alpha-Synuclein Oligomers in Solution

**DOI:** 10.1101/2020.03.10.985804

**Authors:** William E. Arter, Catherine K. Xu, Marta Castellana-Cruz, Therese W. Herling, Georg Krainer, Kadi L. Saar, Janet R. Kumita, Christopher M. Dobson, Tuomas P. J. Knowles

**Author notes:** These authors contributed equally (William E. Arter, Catherine K. Xu).

## Abstract

Oligomeric intermediates are implicated as neurotoxins in the pathogenesis of protein mis-folding diseases. Structural, biophysical and biochemical characterisation of these species is challenging due to their heterogeneous and transient nature, and their typically low abundance. Here, we show that microfluidic free-flow electrophoresis is capable of separating heterogeneous oligomer mixtures on a timescale of seconds, at least two orders of magnitude faster than conventional techniques. This enables analysis of oligomer structural heterogeneity, zeta-potential and immunochemistry with minimal sample perturbation under physiologically-relevant conditions.

## Introduction

Protein misfolding is a molecular hallmark of a number of increasingly prevalent human diseases. ^1^ Amyloid fibrils formed from misfolded proteins are the major components of the Lewy bodies and senile plaques found in the brains of patients with neurodegenerative conditions such as Parkinson’s and Alzheimer’s diseases, respectively.^2^ However, fibrillar species possess low inherent toxicity themselves, and it is instead pre-fibrillar, oligomeric aggregates that are implicated as the principal cytotoxic agents in these disorders. ^3–5^ The characterisation and quantitation of such oligomers in the context of their toxicity and aggregation propensity is therefore an area of intense interest. ^6;7^ However, deciphering the structural attributes of oligomeric species, which exist as an intrinsically heterogenous population of structures during protein aggregation, is not possible using established methods that report the properties of analyte mixtures in an ensemble-averaged manner. Although measurements of oligomer populations, taken throughout protein aggregation processes, have enabled coarse-grained relationships between structure and toxicity to be defined, ^8;9^ they do not permit detailed characterisation of oligomer properties including ζ-potential, which may also play a crucial role in determining oligomer aggregation propensity. ^10^ Furthermore, oligomers exist in a complex, dynamic milieu of protein-protein interactions, but current methodologies are often incapable of probing such systems on a timescale relevant to temporal changes in oligomer populations. ^11^ Crucially, current methods for the separation of heterogeneous protein mixtures such as size exclusion chromatography (SEC), analytical ultra-centrifugation (AUC) and native gel electrophoresis operate on a timescale of minutes to hours, rather than seconds as enabled by the approach shown here. In addition, existing techniques for oligomer analysis often perturb the system in question by dilution^12^ or use of non002Dsolution state approaches, ^13;14^ whereas our method allows arbitrary sample conditions and solution-phase0020analysis.

To address these challenges, we have developed an approach based upon multi-spectral microchip free-flow electrophoresis (µFFE) that achieves rapid solution-phase fractionation and in-situ analysis of heterogeneous mixtures of protein oligomers and monomers. We focus on the aggregation and oligomerisation of alpha-synuclein (αS), a protein that is strongly implicated in the pathogenesis of Parkinson’s disease. ^15^ Using this approach, complex oligomeric mixtures are fractionated on-chip to afford rapid oligomer quantification and characterisation of ensemble heterogeneity, whilst allowing simultaneous measurement of oligomer ζ-potential, a hitherto inaccessible parameter. In addition, the short experimental timescale (analysis time ∼5 s), solution-phase conditions and minimal sample dilution enable an accurate, native-state snapshot of dynamic oligomer populations with high temporal resolution. First, we demonstrate the principle of our method through analysis of stable, fluorophore-labelled kinetically trapped αS oligomers, ^11^ before applying the technique to the analysis of transient oligomers that arise during protein aggregation. Moreover, by use of an oligomer-selective aptamer probe, we further demonstrate the broad applicability of our method by investigation of wild-type, unlabelled αS oligomers.

## Results and Discussion

To begin, we focus on the electrophoretic separation of monomeric αS from a mixture of kinetically-trapped, stable, αS oligomers. ^11^ Our approach is based on a µFFE^16;17^ platform^18^ that allows rapid fractionation of samples containing a complex mixture of oligomeric and monomeric proteins (Figure 1(a)). The sample, flanked by an auxiliary buffer, is passed under laminar flow through the microfluidic chip whilst an electric field is applied perpendicular to the flow direction, resulting in fractionation of the heterogeneous mixture according to the different electrophoretic mobilities of the sample components (Figure 1(b)). Notably, in contrast to other µFFE platforms, our device incorporates an in-line sample-desalting module for rapid (∼2 s) sample preparation on chip (Figure S1, Supporting Information). This enables the use of high-salt, physiologically-relevant buffers such as PBS, which are otherwise inaccessible in µFFE, as excessive ionic conduction through the buffer prevents the application of electric field. We introduce αS monomer labelled with a fluorophore (Alexa546) orthogonal to that of the oligomeric protein mixture (Alexa488) as an in-situ reference, to allow facile in-situ differentiation between monomeric and oligomeric αS fractions (Figure 1(b)). For this approach to be effective, it is necessary for both labelling variants to possess the same electrophoretic mobility; the dyes were chosen due to their same inherent charge, ^19^ and quantification of the mobilities of the two labelling variants confirmed that they were identical, with *µ* = −1.43 ± 0.11 × 10^*−*8^*m*^2^*V* ^*−*1^*s*^*−*1^ (Figure S2).

**Figure 1:**
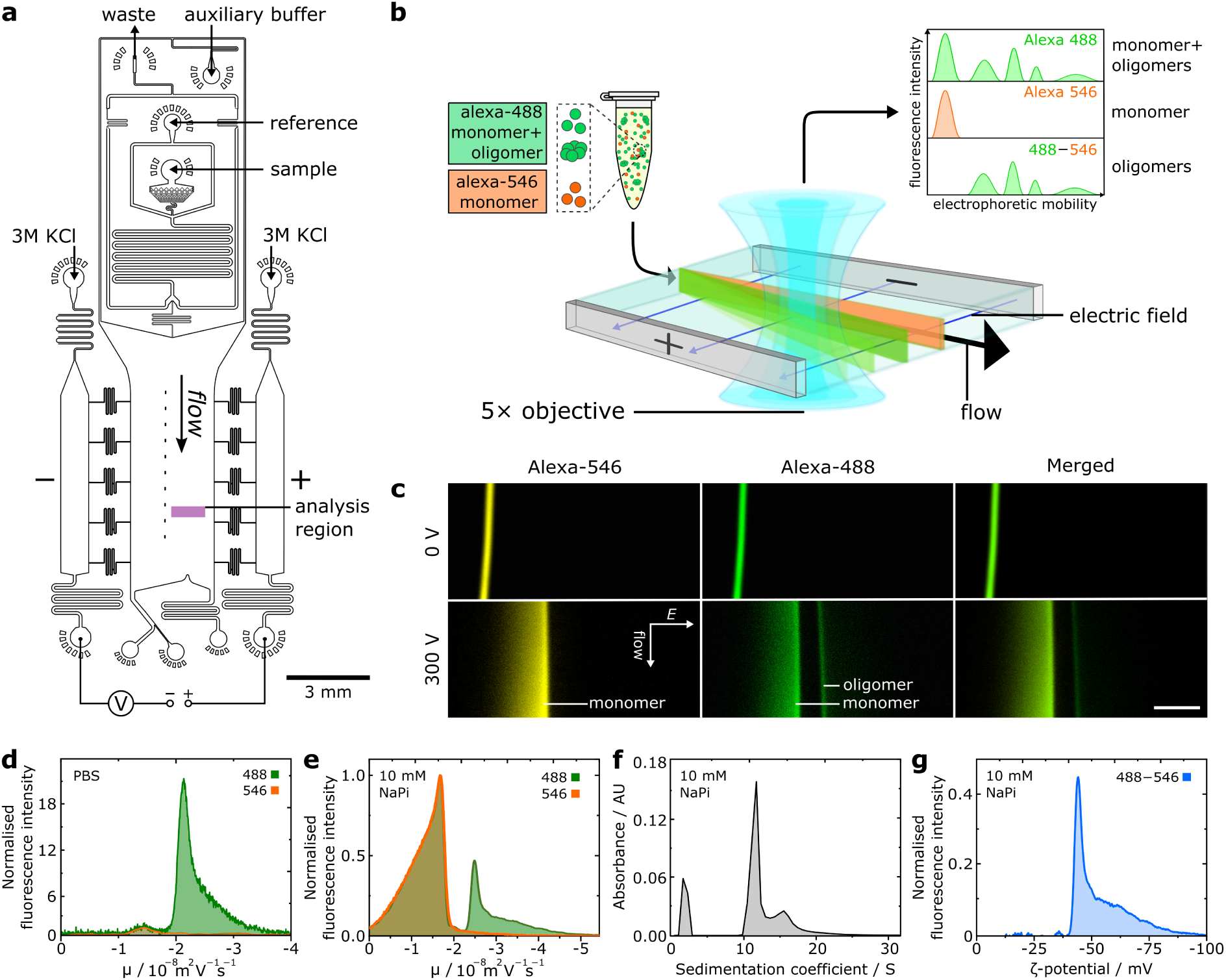
(a) Schematic of device design for desalting-µFFE. The device operates through liquid KCl electrodes, allowing voltage application downstream of the electrophoresis chamber to prevent disruption of flow by electrolysis products. ^18^ Reference sample can be introduced on-chip in pure water diluent, enabling rapid in-situ sample desalting prior to downstream electrophoresis. (b) Schematic for two-colour microfluidic free-flow electrophoresis of oligomeric αS. (c) Fluorescence images of Alexa546-labelled αS monomer reference and Alexa488-labelled oligomeric mixture undergoing µFFE in 10 mM sodium phosphate (NaPi) buffer (pH 7.4). Direction of fluid flow and electric field (*E*) shown by arrows, scale bar = 300 µm. (d) Electropherograms for Alexa546 monomer reference and Alexa488-labelled αS oligomeric mixture fluorescence in PBS buffer. (e) Electropherograms for αS monomer and oligomeric mixture in 10 mM NaPi buffer. (f) AUC data for the same oligomer sample for data shown in (e), the peak at S ≈ 2 is due to monomeric protein. (g) Distribution of oligomer ζ-potential.

We initially demonstrate separation of αS monomers from kinetically-trapped, stable αS oligomers prepared both in high and low-salt conditions (Figure 1(c)). Mixtures of kinetically stable, Alexa488-labelled oligomers together with both Alexa488 and 546-labelled monomer in PBS or 10 mM sodium phosphate buffer were analysed, two distinct fractions were observed for the oligomeric mixture (Alexa488), but only a single fraction was present for the monomer-only sample (Alexa546). By comparison to the Alexa546 fluorescence profile, the lower and higher mobility fractions could be clearly identified as monomeric and oligomeric protein, respectively (Figure 1(d, e)). As expected, a broad peak width is observed for monomer, resulting from faster diffusion due to its smaller hydrodynamic radius in comparison to oligomeric species. This analysis revealed significant oligomer heterogeneity, with a similar electrophoretic profile for both conditions comprising a major oligomer population at *µ* = −2.49±0.16 × 10^*−*8^*m*^2^*V* ^*−*1^*s*^*−*1^ combined with a significant high-mobility shoulder composed of smaller sub-populations. Additional data, definition of apparent mobility and further discussion of desalting-µFFE are provided in the Supporting Information.

We attribute the electrophoretic separation of oligomer and monomer to the faster scaling of effective oligomer charge relative to oligomer size, for an oligomer of *n*_*m*_ monomer units. Oligomer electrophoretic mobility (*µ*_*o*_) is proportional to oligomer charge (*q*_*o*_) and inversely proportional to oligomer hydrodynamic radius (*r*_*o*_) according to 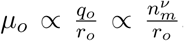, where *ν* is a scaling exponent that links *q*_*o*_ with *n*_*m*_. For spherical oligomers, as in the case of αS,^11^ we approximate that 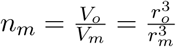 where *V*_*o*_, *V*_*m*_ and *r*_*m*_ represent the oligomer and monomer volumes and monomer hydrodynamic radius, respectively. Together, these expressions yield the oligomer electrophoretic mobility as a function of *n*_*m*_ according to Equation 1, where 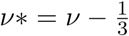. According to this relationship, oligomers are expected to have higher mobilities than monomeric protein, and oligomer mobility is predicted to increase with oligomer size.

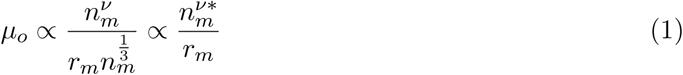

To verify this size-mobility relationship, we compared µFFE to AUC, which is an established technique for generating size distributions of complex samples. A similar oligomer profile was consistently observed in both the AUC and µFFE analyses, indicating a scaling of oligomer electrophoretic mobility with oligomer size (Figure 1(f), additional data and rationalisation are provided in the SI). This observation shows that the electropherograms generated through µFFE correspond directly to size distributions of component species, as predicted by Equation 1, thus validating the µFFE approach as a tool for the analysis of oligomer structural heterogeneity. From the AUC analysis, values of *r*_*o*_ were found to be in the range 5.8-11 nm, in agreement with those determined previously.^11^ Moreover, by comparison of the µFFE and AUC data, it was possible to approximate a value for the scaling exponent *ν* = 0.56 (Supporting Information). *ν* describes the scaling of net oligomer charge with oligomer size according to 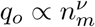 representing a novel and previously inaccessible parameter for oligomer structural characterisation, as *ν* may be sensitive to variation in oligomer quaternary structure due to differences in monomer packing density or coordination, for example.

Having shown that our µFFE approach enables the fractionation of oligomeric protein mixtures, we analysed the resultant oligomer electropherograms to access the distribution of oligomer ζ-potential, a fundamental parameter of nanoscale aggregates. The ζ-potential describes interactions between particles, where it modulates the propensity of the system to aggregate further, and between aggregates and other biological components such as cell membranes. Previously, this parameter has been challenging to study for oligomeric protein aggregates, due to the high degree of structural heterogeneity intrinsic in oligomer samples that confounds conventional techniques, such as dynamic light scattering, for ζ-potential analysis. ^20^ Since our method allows direct observation of oligomer heterogeneity during free-solution electrophoresis, ζ-potentials can be accurately assigned to individual oligomer populations rather than as a population average. Following normalisation, the monomer-only Alexa546 profile was subtracted from the Alexa488 signal to yield an electropherogram corresponding to oligomeric species only, and values for oligomer ζ-potential were extracted from the reported electrophoretic mobilities (Figure 1(e), see SI).^21^ The most common oligomer ζ-potential of ζ = −42.6 ± 4.1 mV is typical of electrostatically stable particulate systems, in agreement with previous studies that have quantified the electrostatic contribution to amyloid formation. ^22;23^ Notably, despite the relationship between surface charge and colloidal stability being of clear relevance to protein aggregation, few experimental studies have attempted to quantify ζ-potential within heterogeneous oligomer populations, likely due to the challenges such an experiment would present for established techniques. ^24^ The µFFE method offers a facile method for quantifying this parameter in protein aggregation systems, introducing an important parameter for understanding the physical-chemical nature of aggregate species.

Following our analysis of kinetically-trapped, stable oligomers, we applied our µFFE platform to the study of transient αS oligomers formed during protein aggregation. Aliquots were periodically withdrawn from the shaking-initiated aggregation reaction of Alexa488-labelled αS monomer, fibrillar components were removed by centrifugation before the samples were analysed by µFFE. High-mobility fractions in the aggregation mixture corresponding to oligomeric αS could be identified and quantified (Figure 2(a, b, c)), following comparison of the electrophoresis profiles for the aggregation mixture and the Alexa546-labelled monomer reference, which yielded electropherograms corresponding to oligomeric αS alone (Figure 2(d)).

**Figure 2:**
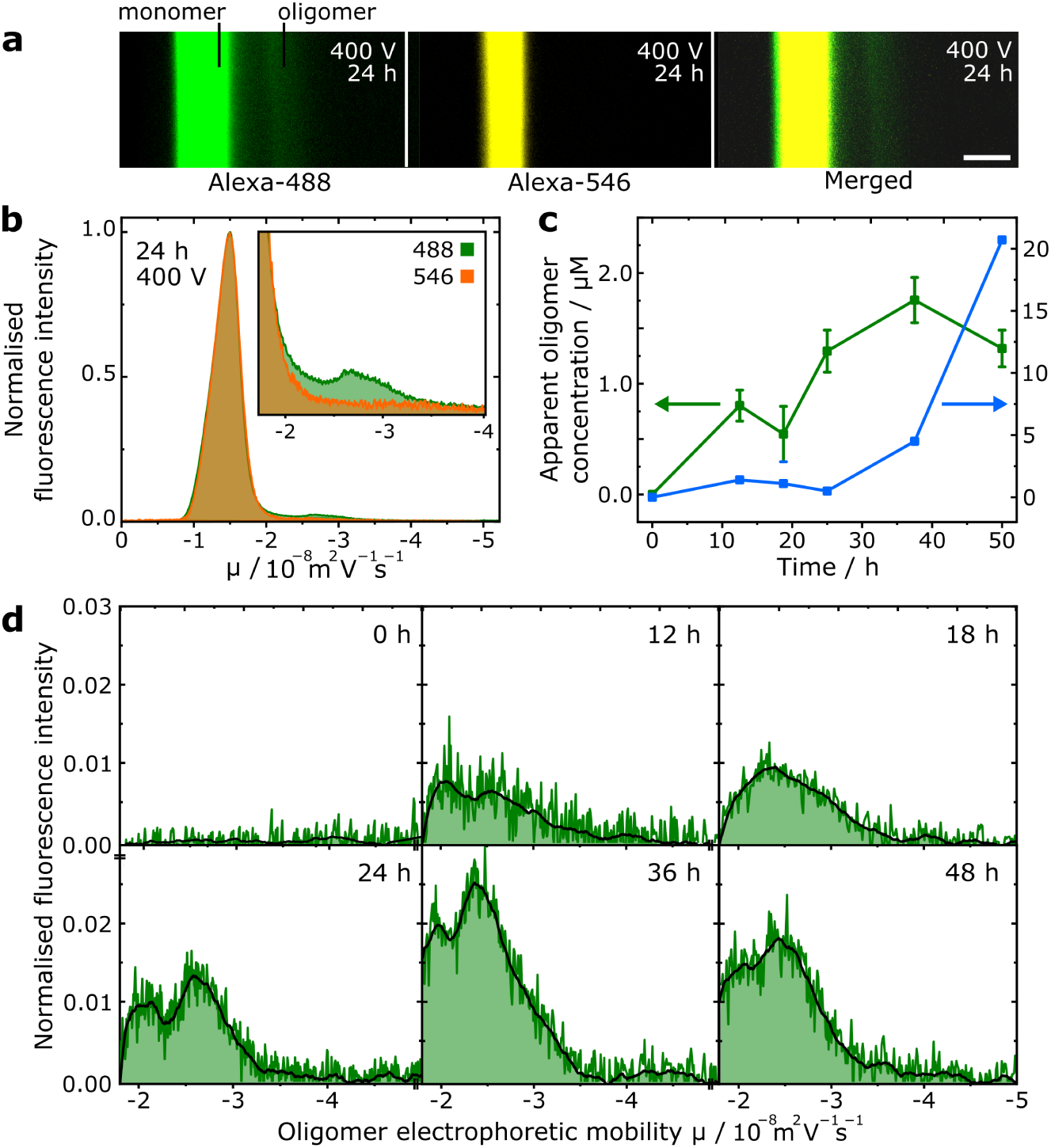
(a) Fluorescence images of µFFE experiment of transient oligomers formed after 24 h αS aggregation. Scale bar = 300 µm. (b) Superimposed electropherograms of normalised alexa-488 and 546 fluorescence for micrographs shown in (a), (inset) magnification of electropherograms showing oligomeric region. (c) Scatter plot showing progression of oligomer concentration (monomer equivalents) and extent of fibril conversion over the sampled time points. Error bars correspond to standard deviation of three repeat measurements of the same sample. (d) Electropherograms and smoothed traces corresponding to oligomeric species only at each sampled aggregation time point.

No oligomers were observed prior to aggregation initiation, before their concentration increased through the lag phase of the reaction, reaching a peak concentration of 1.75 µM (monomer equivalent) with concomitant initiation of protein fibrillisation after 36h. This behaviour is in good agreement with studies employing single-molecule FRET (smFRET) experiments. ^25^ Furthermore, two distinct oligomeric subpopulations are observable in the aggregation timecourse, which corresponds well with smFRET experiments where two major subpopulations of αS oligomers have been observed. ^8;9^ Although our recorded values of oligomer concentration are approximately three times larger than those reported by smFRET techniques, we suggest this is because of inherent under-sampling in the smFRET approach, due to oligomer dissociation caused by sample dilution. ^12;26^ Dilution is not required for µFFE, enabling accurate oligomer quantitation as evidenced by close agreement with values obtained by other techniques such as size exclusion chromatography.^8^ Notably, the µFFE experiment (≈ 5 s analysis time) is three orders of magnitude faster than methods such as AUC. This crucial feature allowed access to transient species that would be otherwise challenging to observe by AUC or other bulk separation methods, with minimal sample consumption of only a few µL.

Further to the fluorophore-tagged protein systems discussed so far, µFFE can be employed for the analysis of wild-type oligomeric species when used in conjunction with an appropriate oligomer-binding probe. We demonstrate this capability by utilising a fluorophore-tagged aptamer selective for αS oligomers. ^27^ As shown previously,^28^ protein–aptamer binding retards the electrophoretic mobility of the aptamer, enabling separation of the bound and unbound fractions (Figure 3(a)). Aliquots taken from stirring-induced aggregation of wild-type αS were incubated with the aptamer and analysed by µFFE (Figure 3(b)). After 4.25 h aggregation time, an additional fraction at lower mobility than the unbound aptamer was observed. By peak integration, an upper bound of 80 nM oligomer concentration could be estimated (Figure 3(c)), indicating, by comparison to labelled oligomer quantitation ((Figure 2(c)), that the aptamer may be selective for a particular subset of the overall oligomer population. The oligomeric nature of the peak was confirmed by µFFE of the aptamer in the presence of αS monomer alone and sonicated αS fibrils (Supporting Information). These data show that label-free, wild-type oligomers can be observed and quantified by µFFE, and demonstrates the versatility of our method towards applications involving immuonchemistry-based oligomer detection and manipulation.

**Figure 3:**
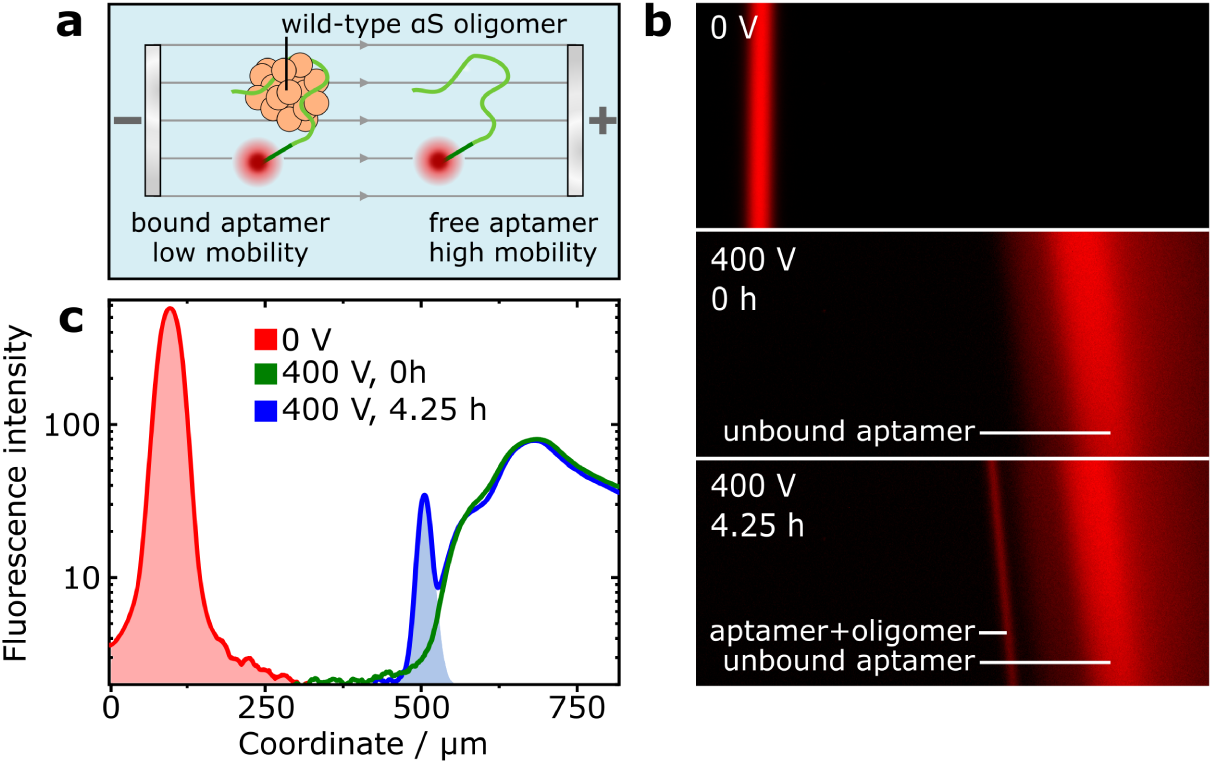
(a) Schematic for fractionation of oligomer-bound and unbound aptamer by µFFE. (b) Images of aptamer fluorescence during µFFE, from aptamer mixed into an aggregation reaction of wild-type αS at 0 h and 4.25 h timepoints. Additional, lower-mobility fraction after 4.25 h indicates aptamer-oligomer binding. (c) Analysis and integration of aptamer electropherograms enable approximation of the concentration of aptamer-bound oligomers.

In conclusion, the µFFE platform presented here is a multifaceted tool for the observation and analysis of oligomeric proteins relevant to misfolding disease. µFFE affords facile, rapid analysis of structural heterogeneity within populations of transient, on-pathway oligomers formed during aggregation under native conditions, with the key advantage that unstable, weakly-interacting oligomers are not perturbed by sample dilution. Electrophoretic analysis enables electrostatic characterisation of oligomers according to their ζ-potential, despite their inherent heterogeneity. Significantly, the µFFE method also facilitates analysis of wild-type, label-free oligomers and their interaction with immuno-reagents. Oligomer-µFFE has many potential applications, such as in the analysis of protein aggregation kinetics, the links between oligomer structure and toxicity, and for probing protein–protein and protein–small molecule interactions relevant towards the design of therapeutic interventions.

## Methods

### Expression and purification of α-synuclein

Wild-type α-synuclein was expressed and purified as previously described. ^29^ Briefly, Escherichia coli BL21 cells overexpressing α-synuclein were collected by centrifugation (20 min, 4000 rpm, 4 °C) in a JLA-8.1000 rotor in a Beckman Avanti J25 centrifuge (Beckman Coulter), resuspended in lysis buffer (10 mM Tris, 1 mM EDTA, protease inhibitor), and lysed by sonication on ice. Following centrifugation (JA-25.5 rotor, 20 min, 18,000 rpm, 4 °C), heat-sensitive proteins were precipitated out of the lysate supernatant by boiling, and subsequently removed by centrifugation (JA-25.5 rotor, 15 min, 18,000 rpm, 4 °C). DNA was precipitated out by incubation with streptomycin sulphate (10 mg/mL, 15 min, 4 °C), and removed by centrifugation. α-synuclein was precipitated out of the supernatant by the slow addition of ammonium sulphate (361 mg/mL) while stirring (30 min, 4 °C). The pellet containing α-synuclein was collected by centrifugation (JA-25.5 rotor, 15 min, 18,000 rpm, 4 °C) and resuspended in 25 mM Tris buffer, pH 7.4, 20 °C. Following dialysis to ensure complete buffer exchange, the protein was loaded onto a HiLoadTM 26/10 Q Sepharose high performance column (GE Healthcare), and eluted at ∼350 mM NaCl, 20 °C with a salt gradient from 0 M to 1.5 M NaCl. Selected fractions were subsequently loaded onto a Superdex 75 26/60 (GE Healthcare) at 20 °C and eluted in PBS, pH 7.4, 20 °C. Protein concentration was determined by absorbance at 275 nm, using an extinction coefficient of 5600 M^-1^cm^-1^. The cysteine-containing variant (N122C) of α-synuclein was purified by the same protocol, with the addition of 3 mM DTT to all buffers.

### Labelling of α-synuclein

α-Synuclein protein was fluorophore-labelled to enable visualisation by fluorescence microscopy. ^11^ In order to remove DTT, cysteine variants of α-synuclein were buffer exchanged into PBS or sodium phosphate buffer by use of P10 desalting columns packed with Sephadex G25 matrix (GE Healthcare). The protein was then incubated with an excess of Alexa-488 or Alexa-546 dyes with maleimide moieties (Thermofisher Scientific) (overnight, 4 °C on a rolling system) at a molar ratio of 1:1.5 (protein-to-dye). The labelling mixture was loaded onto a Superdex 200 16/600 (GE Healthcare) and eluted in PBS or 10 mM sodium phosphate pH 7.4 buffer at 20 °C, to separate the labelled protein from free dye. The concentration of the labelled protein was estimated by the absorbance of the fluorophores, assuming a 1:1 labelling stoichiometry (Alexa-488: 72000 M^-1^ cm^-1^ at 495 nm, Alexa-546: 112 000 M^-1^ cm^-1^ at 556 nm).

### Preparation of stable α-synuclein oligomers

α-synuclein oligomers were prepared by the following procedure. ^11^ Monomeric α-synuclein was lyophilised in Milli-Q water and subsequently resuspended in PBS or 10 mM sodium phosphate buffer, both at pH 7.4, to give a final concentration of ca. 840 uM (12 mg/mL). The resulting solution was passed through a 0.22 uM cut-off filter before incubation at 38 °C for 20-24 h under quiescent conditions. Very small amounts of fibrillar species formed during this time were removed by ultracentrifugation for 1 h at 90000 rpm (Optima TLX Ultracentrifuge, Beckman Coulter, TLA-120.2 Beckman rotor, 288000 g). The excess monomeric protein and some small oligomers were then removed by multiple filtration steps using 100-kDa cut-off membranes. The final concentration of oligomers was estimated on the basis of the absorbance at 495 nm using a molar extinction coefficient of 72000 M^-1^ cm^-1^ for the Alexa-488 labelled oligomers.

### Procedure for generation and isolation of kinetic α-synuclein oligomers

Alexa-488 labelled α-synuclein (N122C, 100 µM, 500-800 µL) was incubated in PBS buffer, pH 7.4, at 37 °C with 0.01% sodium azide, in a 1.5 mL Eppendorf tube with shaking at 200 rpm over 96 hours. 150 µL aliquots were withdrawn and centrifuged for 15 min at 21130 g, to pellet insoluble, fibrillar components of the reaction mixture. The supernatant, containing monomeric and oligomeric α-synuclein, was carefully removed and used immediately for FFE experiments. A small portion of the supernatant was retained for analysis of α-synuclein concentration by UV-vis absorption spectroscopy.

### Microfluidic device fabrication

Devices were designed using AutoCAD software (Autodesk) and photolithographic masks printed on acetate transparencies (Micro Lithography Services). Polydimethylsiloxane (PDMS) devices were produced on SU-8 moulds fabricated *via* photolithographic processes as described elsewhere, ^30;31^ with UV exposure performed with custom-built LED-based apparatus. ^32^ Following development of the moulds, feature heights were verified by profilometer (Dektak, Bruker) and PDMS (Dow Corning, primer and base mixed in 1:10 ratio) applied and degassed before baking at 65 °C overnight. Devices were cut from the moulds and holes for tubing connection (0.75 mm) and electrode insertion (1.5 mm) were created with biopsy punches, the devices were cleaned by application of Scotch tape and sonication in IPA (5 min). After oven drying, devices were bonded to glass slides using an oxygen plasma. Before use, devices were rendered hydrophilic *via* prolonged exposure to oxygen plasma. ^33^

### µFFE device operation

Liquid-electrode µFFE devices were operated as described previously.^18^ Briefly, fluids were introduced to the device by PTFE tubing, 0.012”ID x 0.030”OD (Cole-Parmer) from glass syringes (Gas Tight, Hamilton) driven by syringe pumps (Cetoni neMESYS). µFFE experiments were conducted with auxiliary buffer, electrolyte, monomer reference and sample flow rates of 1200, 250, 140 and 10 µL hr^-1^, respectively, for 15× reduction in buffer salt concentration for samples in PBS buffer.

Potentials were applied by a programmable benchtop power supply (Elektro-Automatik EA-PS 9500-06) *via* bent syringe tips inserted into the electrolyte outlets. Device voltage efficiency was calibrated by comparison of current-voltage curves of the device operating under assay conditions and when filled with 3M KCl electrolyte. Efficiencies were found to be ≈20%, affording electric fields equivalent to 200-267 Vcm^-1^ for potentials of 300–400 V.

Microfluidic experiments were conducted using an inverted fluorescence microscope (Zeiss AxioObserver D1), Alexa488, 546 and 647-labelled species were observed using appropriate filter sets (49002, 49004 and 49006, Chroma Technology) and camera (Evolve 512 CCD, Photometrics).

### Oligomer µFFE experiments

Alexa488-labelled oligomeric mixtures (4 µM monomer equivalent for stabilised oligomers) in either 10 mM sodium phosphate pH 7.4 or PBS buffer were mixed on-chip with Alexa546-labelled monomer (2 µM) in either 10 mM sodium phosphate pH 7.4 or pure water, respectively. For oligomeric samples in 10 mM sodium phosphate or PBS buffer, auxiliary buffer comprised of the same or 15X diluted PBS buffer, respectively, supplemented with 0.05% v/v Tween-20.

Following data acquisition, Alexa488 and Alexa546 fluorescence profiles were extracted, super-imposed and normalised to the peak fluorescence of the Alexa546 monomer. Subtraction of the normalised Alexa546 from the Alexa488 profile afforded profiles due to oligomeric aggregates alone. For quantification of oligomers formed transiently during αS aggregation, the relative peak integrals of oligomeric and monomer populations were compared to the known monomer concentration and degree of sample loss during desalting (see Supporting Information) to calculate the oligomer concentration.

### Aptamer-oligomer µFFE experiments

Wild-type αS (100 µM) was mixed with Alexa647-labelled aptamer (2 µM, Integrated DNA Technologies, sequence is provided in the Supporting Information) in PBS buffer and underwent aggregation by rapid stirring. ^34^ 150 uL aliquots were withdrawn every hour and fibrillar material was removed by centrifugation. The sample then underwent desalting-µFFE analysis as described above.

### Analytical ultra-centrifugation

Sedimentation velocity measurements^35^ were carried out at 20 °C, 43000 rpm (136680 g), using a Beckman Coulter Optima XL-1 analytical ultracentrifuge equipped with UV-visible absorbance optics and an AN50-Ti rotor. The sedimentation coefficient distributions, corrected to standard conditions by using the SEDNTERP programme, were calculated via least-squares boundary modelling of sedimentation velocity data using the c(s) and ls-g*(s) methods, as implemented in the SEDFIT programme. ^36^

## Author Contributions

W.E.A, C.K.X, J.R.K, C.M.D, T.P.J.K designed research, W.E.A, C.K.X performed research, M.C.C, T.W.H, K.L.S contributed reagents/analytic tools, W.E.A, C.K.X analyzed data and W.E.A, C.K.X, J.R.K, T.P.J.K wrote the paper.

## Acknowledgment

The research leading to these results has received funding from the European Research Council under the European Union’s Seventh Framework Programme (FP7/2007-2013) through the ERC grant PhysProt (agreement 337969) and from the Newman Foundation. W. E. A. acknowledges support from the EPSRC Cambridge NanoDTC, EP/L015978/1. C.K.X acknowledges support from a Herchel Smith Research Studentship. T.W.H. acknowledges support from the Oppenheimer Fund and the BBSRC. K.L.S. acknowledges support from the EPSRC. All authors are supported by the Centre for Misfolding Diseases.

