## Supporting Information for "Rapid Fractionation and Characterisation of Alpha-Synuclein Oligomers in Solution"

#### Contents

1 Desalting μFFE 2

2 Validation of two-colour approach 3

3 Resolution in μFFE 5

4 ζ-potential calculations 5

5 Additional analyses of stabilised oligomers 6

6 Analytical ultracentrifugation 7

7 Extraction of $\nu$ from μFFE and AUC data 8

8 Biophysical characterisation of labelled oligomers 8

9 Aggregation of Alexa488-labelled αS 9

10 Aptamer-oligomer interactions 9

### 1 Desalting µFFE

To increase the applicability of our method towards the analysis of oligomeric samples in general, including under physiological conditions, we sought to remove the constraints with regard to buffer type that are a feature of other electrophoresis experiments. µFFE fractionation requires low-salt conditions (e.g. typically $<$ 20 mM NaCl), often facilitated through the use of non-physiological buffering agents,[1] in order to minimise ionic conduction which can otherwise severely limit the electric field that can be induced across the electrophoresis chamber. This is problematic for the analysis of samples in high-salt, physiological buffers such as PBS, and is particularly relevant in the context of protein aggregation given the effect of solution conditions on oligomer stability and formation-dissociation kinetics.[2;3]

A desalting module was incorporated into the µFFE device, upstream of the electrophoresis chamber, in order to rapidly decrease the salt concentration on chip to enable electrophoretic analysis (Figure S1).

The desalting module functions by exploiting the relatively slow rate of diffusion for high molecular weight proteins and protein oligomers in comparison to that of salt ions. Sample solutions undergo diffusional mixing with salt-free water during passage through the desalting module, salts diffuse the full width of the microfluidic channel and are diluted according to the relative flow rates of sample and diluent, whilst proteins only partially diffuse and remain at higher concentration before passage into the electrophoresis chamber. The protein is then re-collected on-chip, before passing into the electrophoresis chamber of the device for µFFE analysis. Although a proportion of the sample is lost using this technique, this effect is minimised for species with slow diffusion such as the large oligomeric complexes of interest here. The degree of sample loss is shown in Figure S1(c, d), with 47% of monomer (r = 3 nm) retained for electrophoresis, increasing to 65–75% for oligomers of r = 6.4–10 nm.[4] Ionic strength can be readily reduced by up to 20×, bringing samples initially in PBS into suitable operating conditions for effective µFFE. Importantly, this desalting process occurs quickly, with the initial desalting step and total time on chip being ∼1.8 and 3.5 seconds, respectively. Therefore, it is predicted that desalting will not significantly affect oligomer structure over the assay timescale, which is significant since solution conditions are known to influence aggregate stability.[2] In addition, the monomer protein reference can be co-injected with the salt-free diluent, minimising the time for potential interactions between reference and analyte.

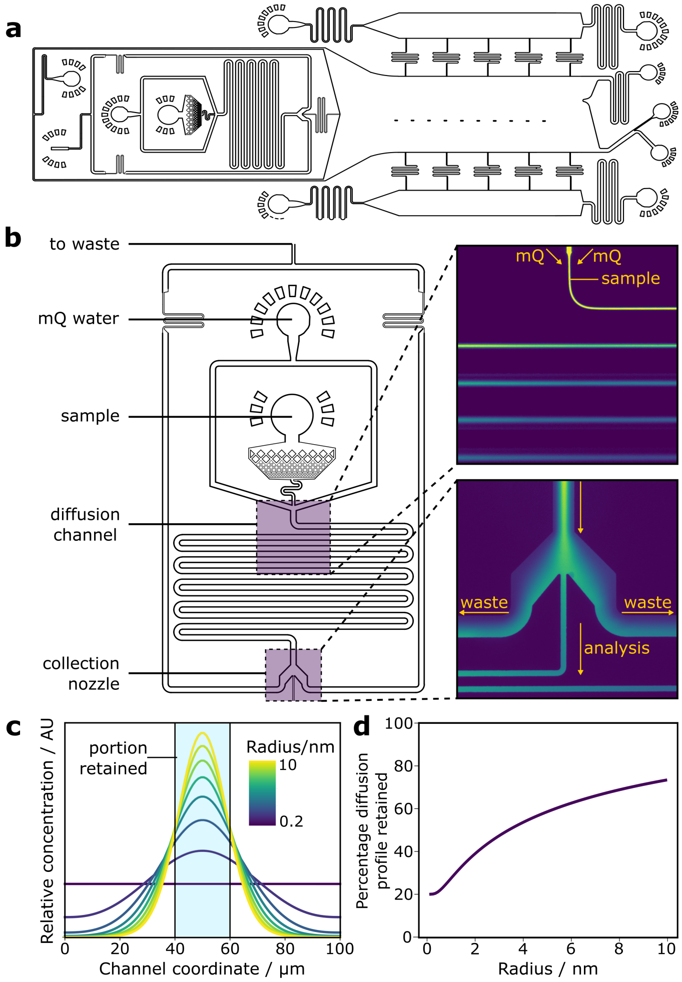


Figure S1: μFFE device with in-line desalting module. (a) Schematic of desalting-μFFE device design. (b) Enlarged schematic of the desalting module. (Inset, upper) Fluorescence micrograph of αS monomer flowing into the desalting-diffusion region. (Inset, lower) Fluorescence micrograph of desalting 'nozzle', showing central portion of αS monomer diffusion profile being selected for downstream μFFE analysis. (c) Schematic showing region of diffusion profiles for r = 0.2 nm ($Na^+$ ions) to r = 10 nm (approximate size of αS oligomers). (d) Plot of percentage of protein retained for downstream analysis as a function of hydrodynamic radius.

#### 2 Validation of two-colour approach

For the dual-colour approach described here, it is necessary that both Alexa488 and Alexa546 labelled αS possess the same electrophoretic properties. Since both dyes possess the same charge ($-2\ e$) and a similar structure, the difference between the labelling variants was expected to be minimal. To verify this prediction, the electrophoretic mobilities of monomeric Alexa488 and Alexa546 labelled αS were quantified using previously described methods (Figure S2).[5] From microscopy images of electrophoretic deflection at the two label wavelengths, no difference could be observed between the two labelling variants (Figure S2(b, c)). To assess this finding quantitatively, the electrophoretic mobilities of the variants were found by plotting the electrophoretic drift velocity as a function of applied electric field. From the linear relationship obtained, it was clear that the measured electrophoretic mobility of $\mu = -1.43 \pm 0.11{\times}10^{-8}\mathrm{m}^2\ \mathrm{V}^{-1}\mathrm{s}^{-1}$ was the same for both fluorophores within experimental error.

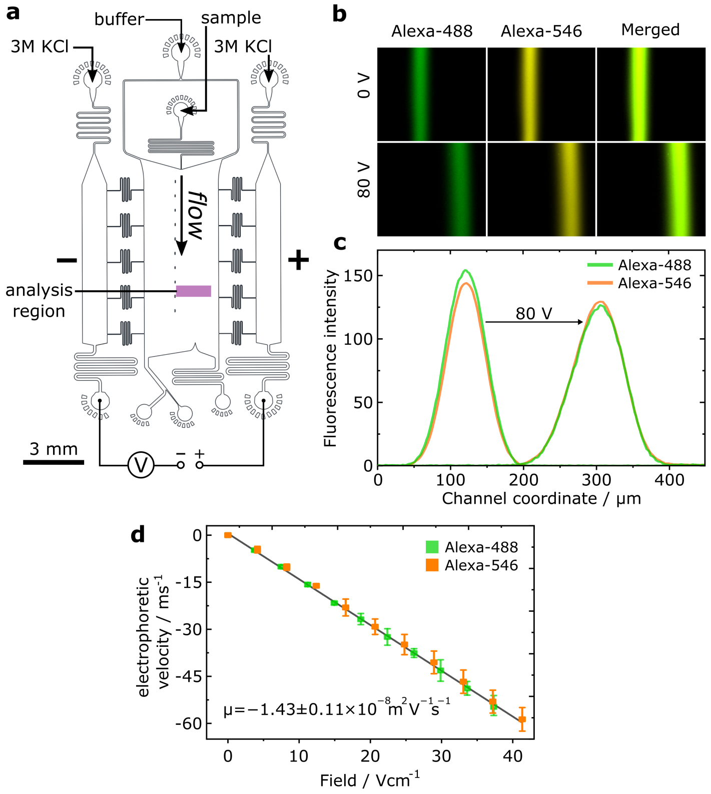


Figure S2: Quantitative FFE of monomeric αS. (a) Design with annotated fluid inlets for the microfluidic FFE device used for quantification of monomeric αS and oligomeric αS in low-salt buffer. (b) Microscopy images of Alexa488, Alexa546 and merged fluorescence of labelled αS monomer, mixed in the same sample and imaged using orthogonal excitation and emission filters, at 0 V and 80 V. (c) Superimposed electropherograms of Alexa488 and Alexa546 labelled αS at 0 V and 80 V corresponding to images shown in (b). (d) Scatter plot and linear fit of Alexa488 and Alexa546 electrophoretic velocities as a function of applied field. Both labelling variants display the same electrophoretic mobility within experimental error.

#### 3 Resolution in μFFE

To account for the finite resolution of the electrophoresis technique, we describe electrophoresis profiles in terms of 'apparent mobility'. Importantly, the mobilities of discrete electropherograms peaks are reported accurately in terms of their absolute value, but there exists an inherent uncertainty for electropherogram positions between intensity maxima. This uncertainty is given by the width of the sample beam under electrophoresis, the variance of which is approximated by:[6]

$$\sigma_{total}^2 = \sigma_{inj}^2 + \sigma_D^2 + \sigma_{HD}^2 = \frac{w_{inj}^2}{12} + \frac{2DL}{v} + \frac{h^2 d^2 v}{105DL}$$

where $\sigma_{inj}^2$, $\sigma_D^2$ and $\sigma_{HD}^2$ describe the injection width, diffusional and hydrodynamic contributions to the total beam width. $w_{inj}$ is the width of the sample injection stream, and $v$, $D$, $L$, $d$, and $h$ are the linear flow velocity, diffusion coefficient, channel length beam deflection and channel height, respectively. For nominal oligomer and monomer radii of 6.4 nm and 3 nm,[7;8] values of $\sigma_{total}$ = 176 μm and 80 μm were calculated for the flow conditions and device geometry used here. Although these values correspond to 28% and 25% of the respective total deflection for oligomer and monomer, an important factor to note is that this band-broadening effect does not prohibit the observation of discrete peaks in oligomer electrophoresis profiles. Thus, the finite resolution of μFFE does not preclude observation and analysis of oligomer heterogeneity.

#### 4 ζ-potential calculations

ζ-potentials are typically calculated for colloidal systems where $\kappa r \gg 1$ using the Smoluchowski relation $\mu = \frac{\epsilon_0 \epsilon_r \zeta}{\eta}$ where $\kappa$ is the inverse Debye length and $r$ is the particle radius. However, using the most common oligomer radius of $r_o = 6.4$ nm as determined by AUC,[4;8] and since $\kappa = 0.52$ nm$^{-1}$ in our experiment (10 mM NaPi), $\kappa r \approx 3.3$ and the Smoluchowski model is therefore an unsuitable model for our data. Instead, the Henry equation (Equation 1):[9]

$$\zeta = \frac{3\mu\eta}{2\epsilon_0\epsilon_r f_H(\kappa a)} \tag{1}$$

provides a better approximation, where $f_H$ is the Henry function, approximated as:[9]

$$f_H(\kappa a) = 1 + \frac{1}{2(1+\delta)^3} \tag{2}$$

$$\delta = \frac{5}{2\kappa a(1 + 2e^{-\kappa a})} \tag{3}$$

from these equations, values of $\delta = 0.7$ and $f_H(\kappa a) = 1.1$ were obtained for $r = 6.4$ nm. Thus a value of $\zeta = -42.6 \pm 4.1$ mV was obtained for the largest population of oligomers where $\mu = -2.49 \pm 0.16 \times 10^{-8}$m$^2$ V$^{-1}$ (maintext, Figure 1(f)).

### 5 Additional analyses of stabilised oligomers

Figure S3 shows additional data for µFFE and AUC analysis of oligomer heterogeneity, from oligmomeric mixtures synthesised in either 10 mM sodium phosphate or PBS buffer. For the NaPi sample, a different oligomer profile is observed compared to the analysis described in the main text, due to differences between samples caused by small variances in oligomer preparation. Importantly, the AUC and µFFE data is self-consistent within each sample dataset, confirming the efficacy of our approach.

In the case of the PBS samples, clear separation between the oligomer and monomer peaks demonstrates effective sample preparation by the microfluidic desalting unit. By comparing the data between the NaPi and PBS-buffered samples, it can be seen that a greater concentration of monomeric αS is present in the lower-salt conditions. This is due to a greater degree of sample dissociation occurring under low ionic strength during the time between oligomer preparation and measurement (2-4 h).[2] Notably, the monomeric fraction is minimal in comparison to the oligomer peak observed in the µFFE data for PBS-buffered oligomers. This implies that no appreciable dissociation of oligomers occurs during the timescale of the µFFE experiment.

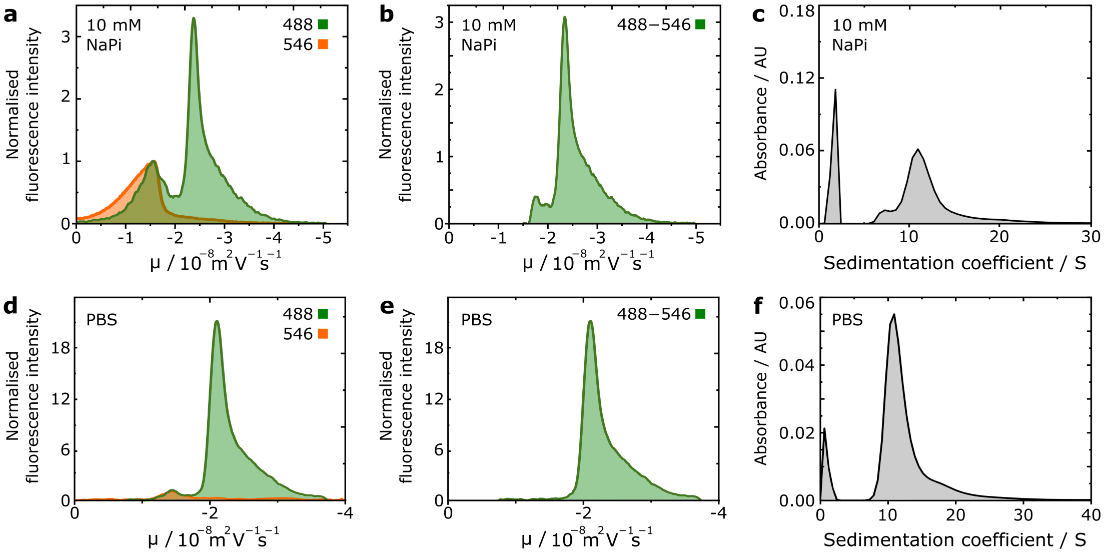


Figure S3: Additional µFFE and AUC data for synthetic oligomers. (a) Superimposed electropherograms for monomer-normalised Alexa546 labelled monomer and Alexa488 labelled oligomer sample in 10 mM NaPi buffer. (b) Electropherogram corresponding to only the oligomeric fraction of an oligomer sample in 10 mM NaPi buffer. (c) AUC data for the same oligomer sample as in (a) and (b), the peak at S ≈ 2 is due to monomeric protein. (d) Superimposed electropherograms for µFFE of monomer-normalised Alexa546 labelled monomer and Alexa488 labelled oligomer sample desalted on chip from PBS buffer. (e) Electropherogram corresponding to only the oligomeric fraction of an oligomer sample desalted on chip from PBS buffer. (f) AUC data for the same oligomer sample as in (d) and (e), the peak at S ≈ 2 is due to monomeric protein.

### 6 Analytical ultracentrifugation

To obtain an independent measure of structural heterogeneity, AUC experiments were conducted using the same two samples of stable αS oligomers. AUC quantifies the sedimentation coefficient ($S$) of particles, defined as the ratio of a particle's sedimentation velocity to the applied acceleration that causes the sedimentation. $S$ is given by Equation 4:

$$s = \frac{m}{6\pi\eta r} \tag{4}$$

where $m$ and $r$ are the mass and radius of the particle. For oligomers, the mass of particles will share a linear relationship with the degree of oligomerisation $n_m$, which is in turn related to the net charge and electrophoretic mobility according to Equations 5-8:

$$\mu_o \propto \frac{q_o}{r_o} \propto \frac{n_m^{\nu}}{r_o} \tag{5}$$

$$n_m = \frac{V_o}{V_m} = \frac{r_o^3}{r_m^3} \tag{6}$$

$$r_o = r_m n_m^{\frac{1}{3}} \tag{7}$$

$$\mu_o \propto \frac{n_m^{\nu}}{r_m n_m^{\frac{1}{3}}} \propto \frac{n_m^{\nu *}}{r_m} \tag{8}$$

Therefore, the sedimentation coefficient may scale with oligomer size in a similar manner to electrophoretic mobility according to Equations 9-10:

$$S_o \propto \frac{m_o}{r_o} \propto \frac{n_m}{r_o} \tag{9}$$

$$\mu_o \propto S_o^{\nu *} \tag{10}$$

where $s_o$ and $m_o$ are the sedimentation coefficient and mass of an oligomer of $n_m$ monomer units, respectively. Therefore, similar observations of oligomer heterogeneity are expected between µFFE and AUC, though these measurements are not directly proportional due to the scaling exponent $0 < \nu* < 1$, that links net oligomer charge to $n_m$.

Oligomer hydrodynamic radii were approximated from AUC data by calculation of oligomer mass as a function of oligomer size, assuming spherical oligomers and protein density[10] $\rho = 1.35$ gcm$^{-1}$ according to:

$$S = 10^{-13} s = \frac{V_o \rho}{6\pi\eta r_o}$$

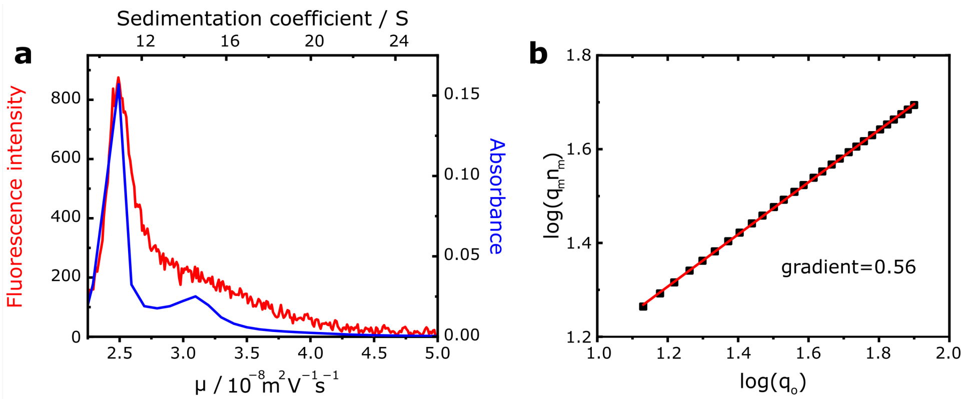


Figure S4: (a) Superposition of µFFE and AUC data for stabilised oligomers in 10 mM NaPi buffer pH 7.4. (b) log-log plot of $q_o$ and $q_m n_m$. Gradient: $\nu = 0.56$.

#### 7 Extraction of $\nu$ from µFFE and AUC data

To approximate the value of $\nu$, which describes the charge-scaling of oligomers according to $q_o \propto n_m^{\nu}$, AUC data was superimposed onto the µFFE electropherogram for stabilised Alexa488-labelled oligomers in 10 mM NaPi buffer pH 7.4 according to the peak and shoulder positions apparent in each dataset (Figure S4(a)). A value of $r_o$ was calculated for each value of $\mu_o$, allowing approximation of $q_o$ as a function of $S_o$. For each value of $S_o$recorded, a corresponding association state in terms of $n_m$ was calculated[4] and the as-determined log values of $n_m$ and $q_o$ were plotted to afford a linear relationship with gradient $\nu = 0.56$.

#### 8 Biophysical characterisation of labelled oligomers

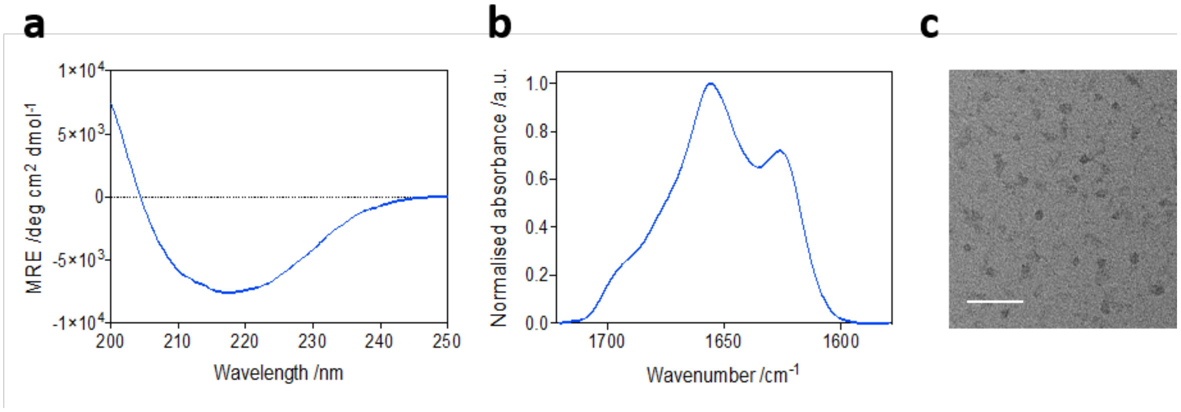


Figure S5: Biophysical characterisation of Alexa488-labelled stable αS oligomers. Representative CD (a) and FTIR (b) spectra, showing the expected antiparallel β-sheet structural content.[4] (c) TEM image of oligomers (scale bar = 100 nm).

#### 9 Aggregation of Alexa488-labelled αS

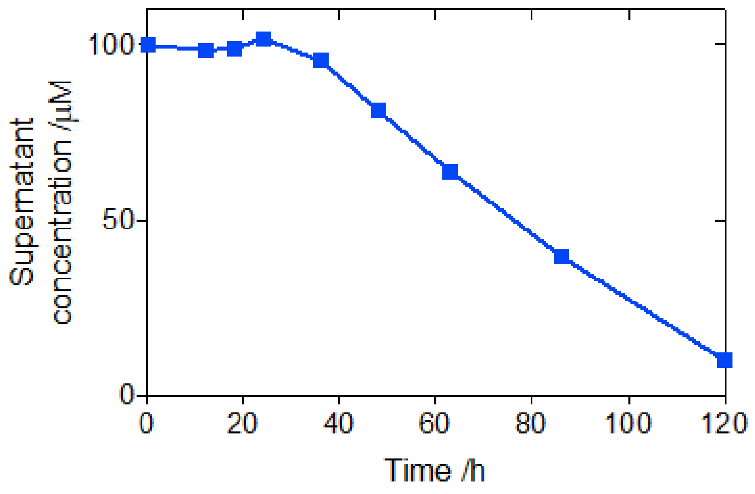


Figure S6: Aggregation kinetics of Alexa488-labelled αS under shaking (200 rpm) at 37 °C, PBS pH 7.4. Fibrils were removed by centrifugation (21,130 rcf, 10 min, 25 °C) and the supernatant concentration quantified by UV-Vis spectroscopy at 495 nm.

#### 10 Aptamer-oligomer interactions

The aptamer used here (Table S1) is known to be strongly selective for oligomers, but also weakly binds fibrillar αS.[11] To exclude the possibility of the lower-mobility band observed in the 4.25 h αS aggregration timepoint being due to aptamer binding to erroneously retained fibrillar material, µFFE was conducted with a sample mixture comprising aptamer and fibrillar αS (Figure S7(a, b)). A clear difference in electrophoretic mobility and electropherogram profile was observed for the aptamer band, in comparison to both the t=0 and t=4.25 h timepoints. This indicates that fibrillar material may also be detectable by µFFE, as suggested previously,[5]and that the band observed at t=4.5 h can be assigned to aptamer-oligomer binding.

Samples of αS fibrils were prepared as described previously.[4] Briefly, monomeric αS at 70 µM in PBS buffer pH 7.4, 0.01% $NaN_3$, was incubated under constant agitation (37 °C, 200 rpm) for 4-6 days. After this time, each sample was centrifuged (15 min at 13200 rpm) and the fibrillar pellet washed twice with PBS before being resuspended into the appropriate volume of PBS. The final concentration of fibrils, that was typically ca. 100 µM in each sample, was estimated by measuring the absorbance at 275 nm using a molar extinction coefficient of 5600 $M^{-1}$ $cm^{-1}$ after disaggregating an aliquot in the presence of 4M guanidinium chloride. To reduce their size to ≈100 nm to prevent blockage of device channels, fibrils were sonicated (20s, 10% power, 30% cycle) prior to use. Aptamer-fibril µFFE was conducted in PBS buffer pH 7.4, with aptamer and fibril (monomer equivalent) concentrations of 2 µM and 100 µM, respectively.

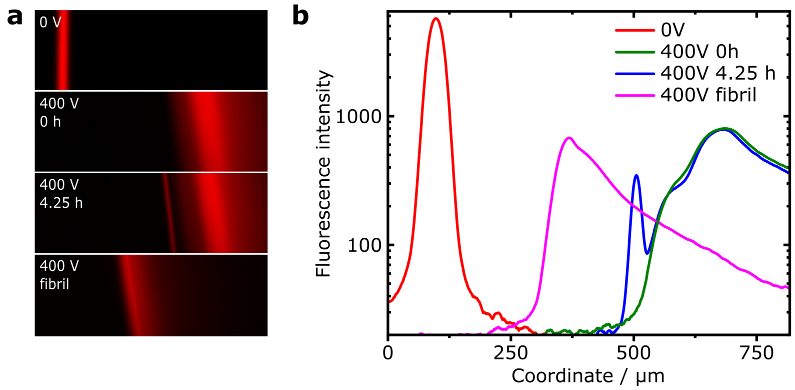


Figure S7: (a, upper three panels) Microscopy images of aptamer fluorescence during µFFE, from aptamer mixed into an aggregation reaction of wild-type αS, as in the main text. (Bottom panel) µFFE of an aptamer-fibril mixture, using sonicated fibrils of wild-type αS. (b) Electrophoresis profiles of aptamer µFFE, as in main text, with addition of profile for aptamer-fibril mixture.

| DNA oligonucleotide | Sequence |
|---|---|
| αS oligomer-selective aptamer | 5'-(Alexa647)-ATTTGCCTGTGGTGTTGGGGCGGGTGCG-3' |

Table S1: DNA sequence for aptamer selective for αS oligomers. Aptameric sequence T-SO508 is underlined.[11] Additional four bases added to provide spacer between aptameric region and fluorophore.
